# Dietary fiber induces a fat preference associated with the gut microbiota

**DOI:** 10.1101/2023.05.30.542811

**Authors:** Yijia Liow, Itsuka Kamimura, Umezaki Masahiro, Wataru Suda, Lena Takayasu

## Abstract

Eating behavior is essential to human health. However, whether future eating behavior is subjected to the conditioning of precedent dietary composition is unknown. This study aimed to investigate the effect of dietary fiber consumption on subsequent nutrient-specific food preferences between palatable high-fat and high-sugar diets and explore its correlation with the gut microbiota. C57BL/6NJcl male mice were subjected to a 2-week dietary intervention and fed either a control (n = 6) or inulin (n = 6) diet. Afterwards, all mice were subjected to a 3-day eating behavioral test to self-select a high-fat or a high-sugar diet. The test diet feed intakes were recorded, and the mice’s fecal samples were analyzed to evaluate the gut microbiota composition. The inulin mice exhibited a preference for a high-fat diet over a high-sugar diet, associated with distinct gut microbiota compositions profiles between the inulin and control mice. The gut microbiota *Bacteroides acidifaciens* (99.68%), *Bacteroides caecemuris* (99.37%), and *Bacteroides xylanolyticus* (92.28%) positively correlated with a preference for fat. Further studies involving fecal microbiota transplantation and eating behavior-related neurotransmitter analyses may clarify the role of gut microbiota on food preferences. Food preferences induced by dietary intervention are a novel observation, and the gut microbiome may be significantly associated with this preference.

## Introduction

Eating is essential to quality health. The study of eating behavior is an interdisciplinary field that has garnered the attention of nutritionists, anthropologists, psychologists, neuroscientists, sociologists, and even economists, to tackle the central question: “*Why do we eat what we eat?* “ Our food choice may seem mundane and arbitrary, yet what we choose to eat has a tremendous effect on our health.

Nutritional scientists have illuminated the physiological and metabolic responses elicited by ingested foods’ quantity and composition [1, 2]. Neuroscientists have targeted brain chemistry and neural pathways that control eating behaviors at the core of human behavior [3–5]. Sociologists and psychologists have developed several models to break down and construct a holistic human food selection framework. Anthropologists have shed light on culture’s imprinted effect on food acquisition, preparation, attitudes, and rituals. Each discipline dives into the key question around eating formulated in their own right; however, dysfunctional eating behaviors are increasing drastically in contemporary post-industrial societies [6, 7], unprecedentedly threatening human health.

In nutritional science, two facets of eating behavior, food intake [8] and preference [9–12], have been extensively studied. Food intake is the quantitative measure of ingested food; common parameters include meal size, frequency, and appetite. Food intake regulation has been in the spotlight because obesity, one of the leading causes of deteriorating quality of life, is induced by a chronic positive energy balance. Dietary fiber is one of the most well-studied models for improving food intake to mitigate obesity [13]. Dietary fiber consumption regulates food intake through two mechanisms: (1) increases satiation and satiety in the host through its physical characteristics [14], and (2) actions of gut-derived hormones on the appetite center in the brain to control food and energy intake [15, 16]. A robust association between dietary fiber intake and eating behavior via the gut-brain axis has been discovered with the recent bloom of gut microbiome studies. Dietary fiber ingestion stimulates the growth of a specific subset of gut microbiota and increases short-chain fatty acid and appetite-related hormone levels [14, 17]. These hormones travel from the gut to the brain and act on the hypothalamus to signal appropriate homeostatic ingestive behavior in the host [16, 18].

Food preference, like food intake, is another key factor in constructing a framework to understand the complexity of human eating behavior. Nutritionists and psychologists have developed instruments to assess food preferences and underlying food motives in human subjects, including the Leeds Food Preference [19] and the Food Choice Questionnaire [20], used across different cultural contexts. Neuroscientists have also conducted extensive interventional studies in animals to examine the neural patterns that serve as the basis of decision-making regarding food, specifically food preferences. The central peptide administration, including neuropeptide Y and opiate in the hypothalamus, elicited a preference for carbohydrate and palatable food consumption [21]. Hormones, including glucagon and growth hormone-releasing hormones, increased protein intake when the mice were offered different macronutrient sources [22–25]. The neurotransmitter serotonin (5-HT) suppresses appetite in high-carbohydrate foods, whereas galanin consistently elicits a preference for high-fat foods [26, 27]. The gut microbiota is responsible for more than 90% of serotonin biosynthesis, a molecule associated with nutrient-specific food preference [28].

Studies targeting the direct and indirect food intake control through dietary fiber and gut microbiota have greatly expanded our understanding of eating behavior. However, to the best of our knowledge, no studies have explicitly addressed dietary fiber intake’s impact on food preferences associated with the gut microbiota. Anatomical and physiological similarities between humans and mice make them suitable models for dietary intervention studies. Our study aimed to use a mouse model to clarify dietary fiber consumption’s physiological effects on food preference between palatable high-fat and high-sugar diets in mice associated with gut microbiota.

## Materials and methods

### Animals and housing

Twelve 8-week-old specific pathogen-free (SPF) inbred C57BL/6NJcl male mice were obtained from CLEA Japan, Inc., Tokyo, Japan. The mice were housed in groups of three, except during the eating behavioral test at the RIKEN Yokohama Campus1^*°*^C and humidity at 50 ± 5% in the SPF animal facility. Animal care and treatment Animal Facility. They were subjected to a 1-week habituation period and fed a control diet, ad libitum (D21052808, Research Diets, Inc., New Brunswick, NJ, USA). All mice were randomly divided into two groups (inulin or control); baseline body weights were measured to ensure the absence of outliers in each experimental group. Ear punching was performed for mouse identification at the beginning of the habituation period. The lights were set to a 12-h light-dark schedule with lights on at 7 a.m. [Zeitgeber (ZT) 0] and off at 7 p.m. (ZT12). The temperature was maintained at 24±1^*°*^C and humidity at 50 ± 5% in the SPF animal facility. Animal care and treatment were conducted according to the institutional guidelines of RIKEN Yokohama Campus. The Ethics Committee of the RIKEN Yokohama campus approved all experimental procedures [Y-H29-170187] and complied with the ARRIVE guidelines.

### Dietary intervention

At 9 weeks old, all experimental mice were divided into two groups (n=6/group) and assigned to one of the following feeding regimes for 2 weeks: (a) high-fiber inulin diet (D21052809, Research Diets, Inc., New Brunswick, NJ, USA) and (b) control diet (D21052808, Research Diets, Inc., New Brunswick, NJ, USA), as shown in Fig 1. The control diet used the AIN-93M as a base with 1% cellulose added. The treatment diet contained approximately the same composition as the control, with 10% inulin added. The control diet comprised 15% protein, 76% carbohydrate, and 4% protein, with an energy density of 4 kcal/g. The high-fiber inulin diet comprised 14% protein, 72% carbohydrate, and 4% fat, with an energy density of 3.88 kcal/g. Both diets’ protein ratio comprised casein (mineral acid 30 mesh) and L-cysteine; the carbohydrate comprised corn starch, maltodextrin 10, sucrose, cellulose, and inulin; the fat comprised soybean oil and t-butylhydroquinone. Table 1 shows a description of the nutritional compositions of the control and treatment diets. The mice’s body weights were measured at the start and end of the dietary intervention.

**Table 1.** Intervention diets nutritional composition

|  | Control Diet | Inulin diet |
| --- | --- | --- |
| Macronutrient (%) |  |  |
| Protein | 15.00 | 14.00 |
| Carbohydrate | 76.00 | 72.00 |
| Fat | 4.00 | 4.00 |
| Energy (kcal) | 4.00 | 3.88 |
| Ingredients (gm/3850 kcal) |  |  |
| Casein | 0.00 | 0.00 |
| Casein, Mineral Acid 30 Mesh | 140.00 | 140.00 |
| L-Cystein | 1.80 | 1.80 |
| Corn Starch | 496.00 | 476.94 |
| Maltodextrin 10 | 125.00 | 125.00 |
| Sucrose | 100.00 | 100.00 |
| Cellulose | 10.00 | 10.00 |
| Inulin | 0.00 | 50.00 |
| Soybean Oil | 40.00 | 40.00 |
| t-Butylhydroquinone | 0.008 | 0.008 |
| Mineral Mix S1022M | 35.00 | 35.00 |
| Vitamin Mix V10037 | 10.00 | 10.00 |
| Choline Bitartate | 2.50 | 2.50 |
| Source: Research Diets, INC (New Brunswick, NJ) |  |  |

**Fig 1.**
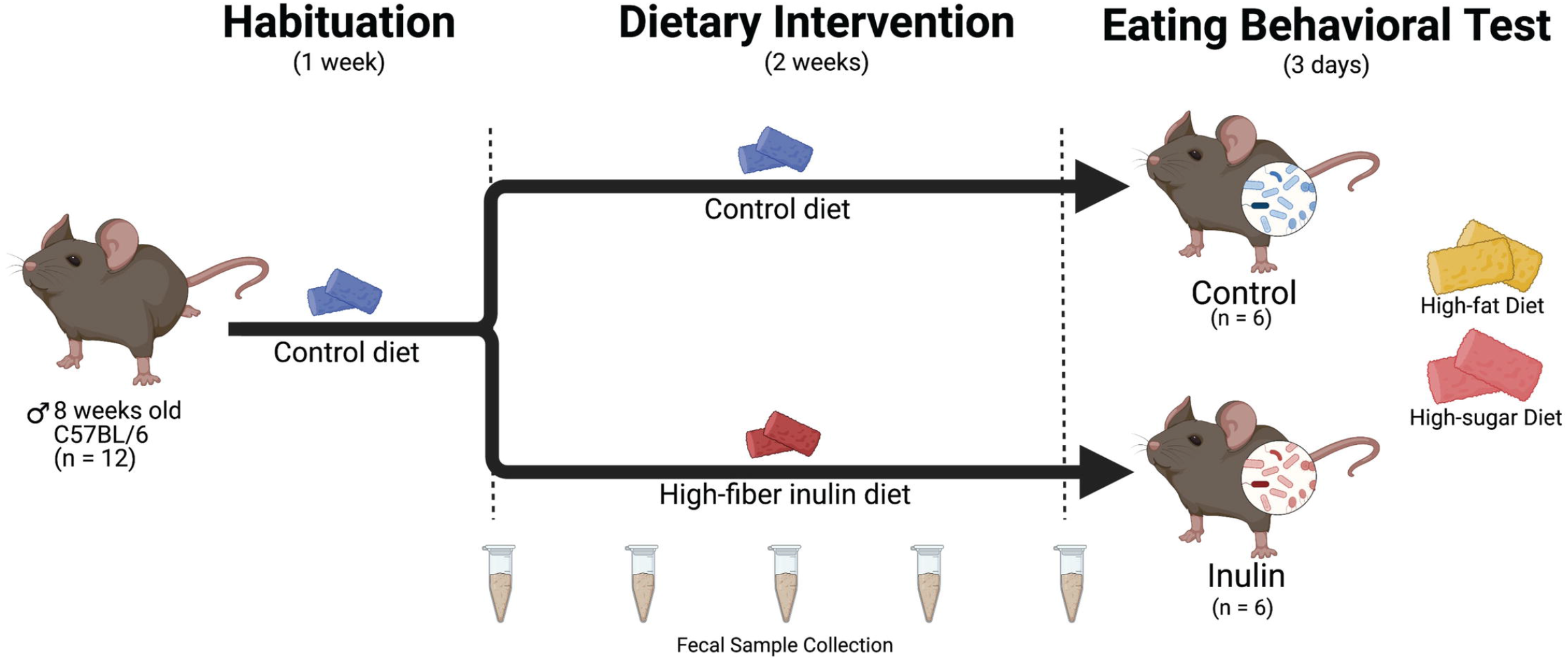
Experimental design overview. Twelve mice underwent a 1-week habituation period followed by a 2-week dietary intervention and were fed one of the following diets: (1) control; (2) high-fiber inulin. After the dietary intervention, the mice were subjected to a 3-day eating behavioral test to choose between palatable high-fat and high-sugar diets. PD, postnatal day. Created with BioRender.com.

### Eating behavioral test

After the 2-week dietary intervention, all experimental mice were housed individually in a customized testing cage (S1 Fig) for a 3-day eating behavioral test to self-select between a palatable high-fat or a high-sugar diet. The testing cage had a 380 × 120 × 115 mm dimension and was made of acrylic materials with three main compartments: two feeding chambers on the left- and right-hand sides and a resting chamber in the middle connecting the feeding chambers. Stainless-steel feeding cages were placed on top of the feeding chambers, where the mice could reach the test diets. A stainless-steel lid was placed above the resting area to prevent the mice from escaping and as support for the water bottle. The test diets were provided by CLEA Japan, Inc. The high-fat diet comprised 14.9% protein, 44.7% carbohydrate, and 40.4% fat, with an energy density of 4.59 kcal/g. Cocoa butter (20%) was the main fat source. The high-sugar diet comprised 17.4% protein, 70.8% carbohydrate, and 11.8% fat, with an energy density of 4.03 kcal/g. Sucrose was the main sugar source (70%). The test diets’ complete nutritional compositions are listed in Table 2. Their positions were switched every 12 h to prevent a location bias. The eating behavioral test was conducted during the dark phase, beginning at 7 p.m. and terminating at 7 a.m. the next day. The behavioral experiment was carried out in a soundproof room where the temperature was maintained at 24 ± 1^*°*^C and humidity at 50 ± 5%. The high-fat and high-sugar test diet feed intake were recorded at the beginning and end of each 12-h test session. In addition to the test diet feed intake, the fat preference score was calculated as follows:

**Table 2.** Test diets nutritional composition

|  | High-sugar diet (HS) | High-fat Diet (HF) |
| --- | --- | --- |
| Macronutrient (%) |  |  |
| Protein | 17.39 | 14.92 |
| Carbohydrate | 70.79 | 44.69 |
| Fat | 11.82 | 40.38 |
| Energy (kcal) | 4.03 | 4.59 |
| Ingredients (g) |  |  |
| Sucrose | 70.00 | 34.90 |
| Cocoa Butter | 0.00 | 20.00 |
| Milk Casein | 20.00 | 19.50 |
| Corn Oil | 0.00 | 0.00 |
| Corn Starch | 0.00 | 15.00 |
| Crystalline Cellulose | 5.00 | 5.00 |
| Mineral Mix (AIN-76) | 3.50 | 3.50 |
| Vitamin Mix (AIN-76) | 1.00 | 1.00 |
| CaCO3 | 0.00 | 0.40 |
| DL-Methionine | 0.30 | 0.30 |
| Choline Deltartate | 0.20 | 0.20 |
| Cholesterol | 0.00 | 0.20 |
| 3-Butylhydroquinone | 0.00 | 0.004 |
| Source: CLEA Japan, INC. |  |  |

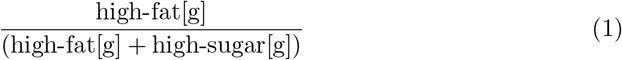

The mice were sacrificed by cervical dislocation at the end of the behavioral test.

### Fecal sample collection and bacterial DNA extraction

Fecal samples were collected before, during, and after the dietary intervention to examine the gut microbiota composition changes induced by dietary fiber inulin. Fresh fecal samples were immediately frozen in liquid nitrogen and stored at -80^*°*^C until further processing. Bacterial pellets were suspended in TE20 and incubated with 15 mg/mL lysozyme (Sigma-Aldrich Co., LCC., St. Louis, MO, USA) and purified achromopeptidase (Wako Pure Chemical Industries, Osaka, Japan) at a final concentration of 100 units/uL at 37^*°*^C for 2 h. Furthermore, 1% (w/v) sodium dodecyl sulfate and 1 mg/mL proteinase K (Merck Millipore, Darmstadt, Germany) were added to the fecal pellets and incubated at 55^*°*^C for 1 h. The lysate was treated with phenol/chloroform/isoamyl alcohol (NIPPON GENE, Tokyo, Japan). Bacterial DNA was precipitated using 3 M sodium acetate and isopropanol, centrifuged at 12,000 g at 4^*°*^C for 5 min, and rinsed with 75% ethanol. RNase (DNAse-free) solution at a final concentration of X was added to the bacterial DNA and incubated at 37^*°*^C for 30 min, followed by 20% PEG6000-2.5 M NaCl to precipitate high-molecular-weight DNA by centrifugation at 12,000 g at 4^*°*^C for 5 min. The final bacterial pellet was rinsed with 75% ethanol three times to remove residual PEG and NaCl, dried under vacuum, and dissolved in 10 mM Tris-HCl/1 mM EDTA.

### 16S rRNA gene Amplicon sequencing

The 16S ribosomal RNA (rRNA) gene V1-V2 region was polymerase chain reaction (PCR)-amplified from the bacterial DNA, using the 16S metagenomic sequencing library protocol (Illumina, Inc., San Diego, CA, USA), and amplified using PCR with universal primers 27F-mod (5’-AGRGTTTGATYMTGGCTCAG-3’) and 338R (5’-TGCTCCTCCCGTAGGAGT-3’). A solution of 44 uL PCR mixture, 2 uL 16S amplicon PCR forward (1 uM) and reverse primers (1 uM), 4 uL bacterial DNA, and PCR-grade water was prepared to a final volume of 50 uL. PCR amplification was conducted with pre-denaturation at 95^*°*^C for 3 min, followed by 20 cycles of 95^*°*^C for 30 s, 55^*°*^C for 30 s, 72^*°*^C for 30 s, and a final extension at 72^*°*^C for 3 min. The PCR products were purified using AMPure XP beads. Purified products were sequenced using an Illumina MiSeq System (Illumina Inc., San Diego, CA, USA). Operational taxonomic units (OTUs) were selected and binned with 97% sequence similarity. Alpha diversity was calculated based on Chao1 and Shannon indices, and beta diversity was clustered using UniFrac principal coordinate analysis (PCoA).

### Statistical analysis

All statistical tests and data visualizations were performed with R version 4.0.2 and RStudio version 1.3.1093. The Wilcoxon rank-sum test was used to test for statistical differences in (1) the high-fat and high-sugar test diet feed intake, (2) the high-fat and high-sugar test diet energy intake, and (3) body weight changes among the inulin and control mice. Permutational multivariate analysis of variance (PERMANOVA) was performed to detect community-level differences between groups and was corrected using the Benjamini-Hochberg method for false discovery.

## Results

### Dietary fiber consumption and nutrient-specific food selection

Fig 2 shows the comparison between the high-fat and high-sugar test diet intakes, including the total feed intake between the two mice groups. Fig 2a shows that the inulin mice significantly preferred the high-fat (P=0.0043) over the high-sugar diet, whereas no significant difference in preference was detected in the control mice (P=0.39). Moreover, high-fat and high-sugar diet intake was compared between the two groups. Inulin mice consumed significantly less high-sugar diet than the control (P=0.041); however, a marginally significant difference in high-fat diet intake was detected (P=0.065) between the two groups. Considering the drastic difference in the test diets’ macronutrient proportions, the inulin and control mice’s combined feed intakes (high-fat + high-sugar) were compared. However, no significant differences in total feed intake were detected (P=0.87), as shown in Fig 2b. We also computed the individual and average fat preference scores in the inulin and control mice (S2 Fig). Inulin mice exhibited a significantly higher fat preference score than the control (P=0.005). These results show that inulin mice prefer a high-fat diet with a suppressed appetite for a high-sugar diet. Furthermore, the high-fat and high-sugar diets’, separately, and total energy intakes were compared between groups, with no significant differences detected (S3 Fig). The inulin and control mice’s body weights increased significantly after dietary intervention (P=0.0026 and 0.0045, respectively, (S4 Fig)). However, no significant differences in body weight after dietary intervention were detected between the two groups (P=0.229).

**Fig 2.**
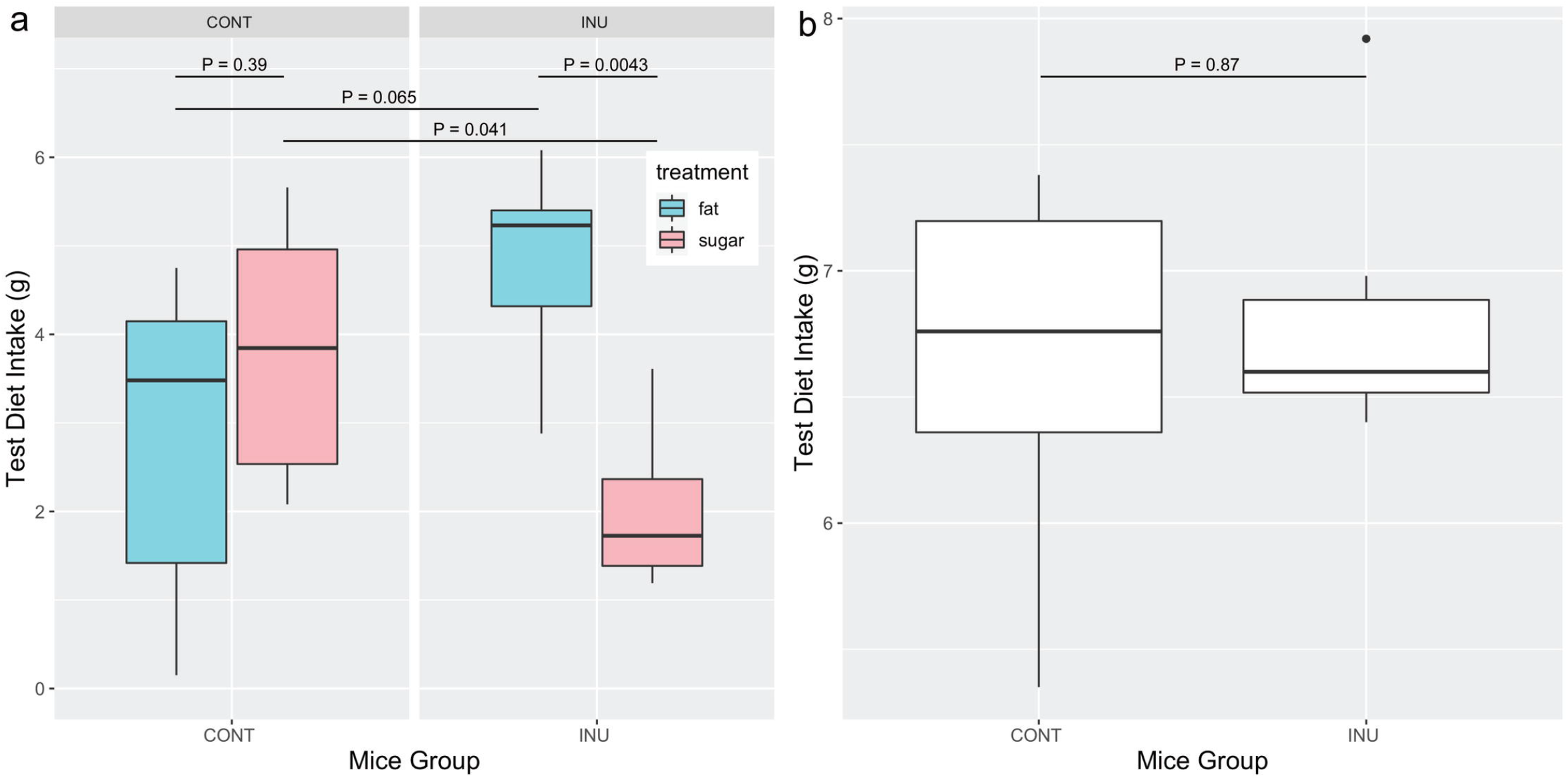
Nutrient-specific food selection results. Fig 2a shows a significant difference in the high-fat and high-sugar feed intakes in Inulin mice (P=0.0043); however, no significant difference was detected in the control (P=0.39). In addition, inulin mice consumed significantly less high-sugar diet than the control (P=0.041); however, no significant difference was detected in the high-fat diet between the two groups (P=0.065). Fig 2b shows no significant differences in the total test diet intake between the two groups (P=0.87).

### Taxonomical features of gut microbiota post-dietary intervention

Gut microbiota analyses were conducted at the genus- and OTU-levels with an abundance of less than 0.5% between the inulin and control mice after dietary intervention on day 14 (Fig 3). Taxonomical assessment combined with the Wilcoxon rank-sum test revealed distinct gut microbiota compositions between the inulin and control mice. Fig 3a illustrates the genus-level taxonomy. The top three most abundant OTUs in inulin mice were *Bacteroides, Lactobacillus*, and *Faecalibaculum*. However, the top three abundant OTUs in the control mice were *Faecalibaculum, Bifidobacterium*, and *Bacteroides*. Fig 3b illustrates the OTU-level taxonomy. The top 10 enriched OTU observed in inulin mice were *OTU00002 Bacteroides acidifaciens* (99.68% identity similarity), *OTU00001 Lactobacillus taiwanensis* (99.71%), *OTU00003 Faecalibaculum rodentium* (99.67%), *OTU00006 Butyricoccus faecihomonis* (81.33%), *OTU00014 Lactobacillus murinus* (100%), *OTU00040 Lactobacillus reuteri* (99.42%), *OTU00004 Bifidobacterium pseudolongum* (99.68%), *OTU00038 Frisingicoccus caecimuris* (89.51%), *OTU00036 Magnetovibrio blakemorei* (83.57%), and *OTU00017 Bacteroides caecimuris* (99.37%). However, the top 10 most abundant OTU observed in control mice were *OTU00003 Faecalibaculum rodentium* (99.67%), *OTU00002 Bacteroides acidifaciens* (99.68%), *OTU00001 Lactobacillus taiwanensis* (99.71%), *OTU00004 Bifidobacterium pseudolongum* (99.68%), *OTU00005 Bacteroides vulgatus* (93.02%), *OTU00025 Anaeromassilibacillus senegalensis* (83.23%), *OTU00007 Prevotellamassilia timonensis* (87.9%), *OTU00013 Eisenbergiella massiliensis* (88.85%), *OTU00010 Gabonia massiliensis* (82.75%), and *OTU00014 Lactobacillus murinus* (100%). A heatmap based on hierarchical clustering was generated to visualize the bacterial composition dynamics after dietary intervention (S5 Fig). Of the 136 OTUs detected in the fecal samples, 47 significantly differed in their relative abundance between the two groups.

**Fig 3.**
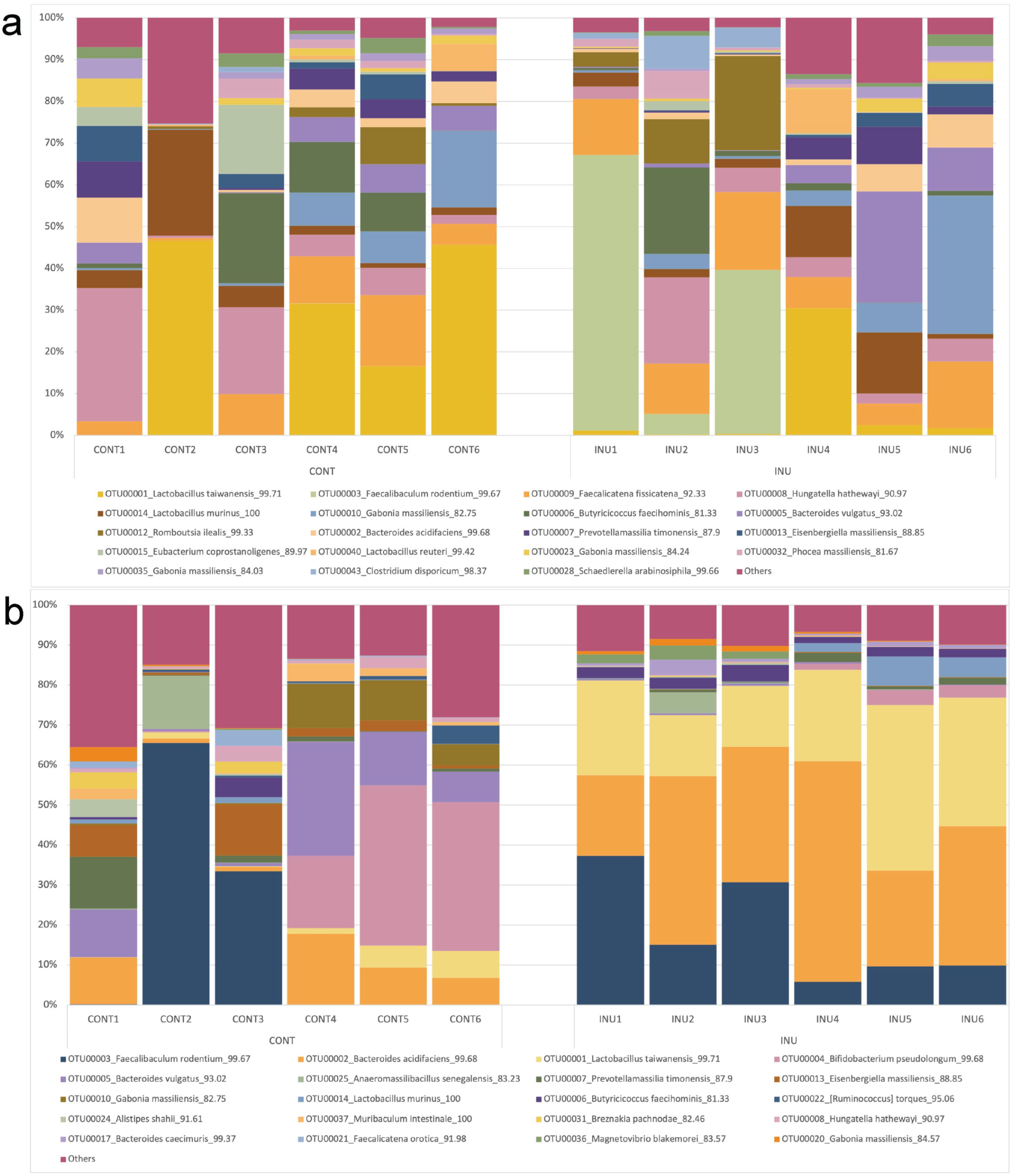
OTU-level taxonomy tables from 16S gene amplicon sequencing of mice fecal samples. Fig 3a: Gut microbial composition of control and inulin mice pre-dietary intervention (D0); Fig 3b: Gut microbial composition of control and inulin mice post-dietary intervention (D14).

PERMANOVA (permutational multivariate analysis of variance) of the weighted UniFrac distances revealed significant structural differences in the gut microbiota composition in the inulin and control mice induced by dietary fiber consumption (S6 Fig). The weighted PCoA UniFrac distance plot considers certain bacterial species’ presence/absence and relative abundance. In the weighted UniFrac distances, PCoA1 captured 28.96% of the total variation from the input data, and PCoA2 captured 27.41% of the total variation in the input data. The PERMANOVA test revealed that significant changes (P=0.009) in inulin mice’s gut microbiota community structure occurred at the third sampling time point during the 2-week dietary intervention.

### Correlation between gut microbiota and nutrient-specific food preference

Fig 4 reveals that 15 OTUs were significantly correlated with hedonic eating behavior, with a preference for fat. Three out of 15 OTUs were positively correlated with fat preference: *OTU00002 Bacteroides acidifaciens* (99.68%), *OTU00017 Bacteroides caecimuris* (99.37%), and *OTU00066 Bacteroides xylanolyticus* (92.28%) (P=0.0059, 0.024, and 0.028, respectively). Conversely, OTUs that were negatively correlated with fat preference were found to be positively correlated with sugar preference, they were: *OTU00008 Hungatella hathewayi* (90.97%), *OTU00021 Faecalicatena orotica* (91.98%), *OTU00026 Anaeromassilibacillus senegalensis* (82.64%), *OTU00029 Fusicatenibacter saccharivorans* (86.29%), *OTU00009 Facelicatena fissicatena* (92.33%), *OTU00071 Anaerocolumna cellulosilytica* (85.94%), *OTU00059 Anaerotignum lactatifermentans* (85.94%), *OTU00034 Colidextribacter massiliensis* (86.69%), *OTU00049 Eubacterium eligens* (89.72%), *OTU00034 Clostridium clostridioforme* (93.52%), *OTU00080 Dorea formicigenerans* (93.65%), and *OTU00164 Gabonia massiliensis* (82.69%) (P=0.033, 0.018, 0.0093, 0.0035, 0.016, 0.023, 0.032, 0.021, 0.038, 0.00018, 0.013, and 0.021, respectively).

**Fig 4.**
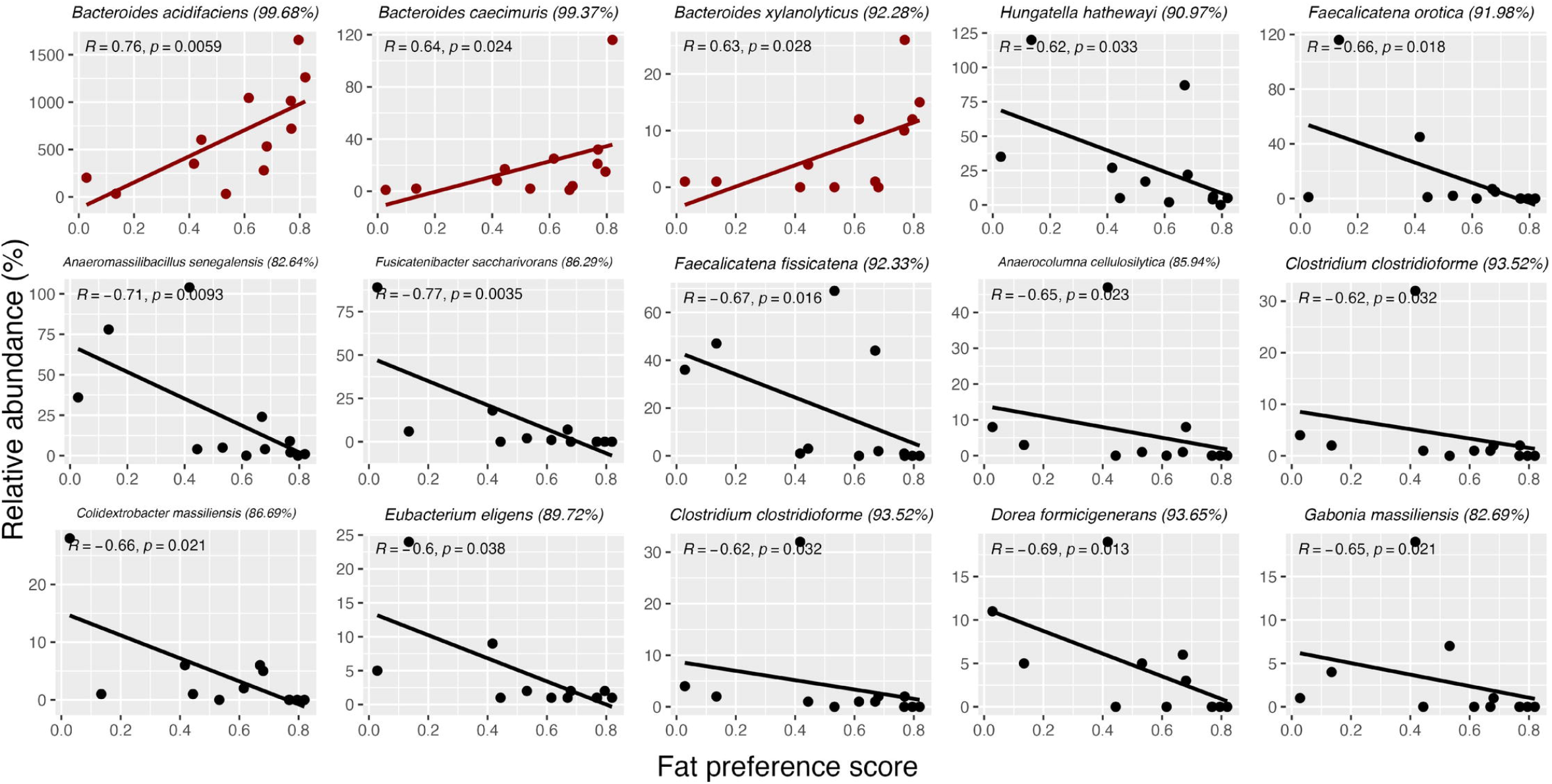
Correlation between fat preference and gut microbiota. Fifteen OTUs significantly correlated with fat preference. Three out of 15 OTUs positively correlated with fat preference: *Bacteroides acidifaciens, Bacteroides caecimuris*, and *Bacteroides xylanolyticus* (P=0.0059, 0.024, and 0.028, respectively).

## Discussion

Food preference is an essential component to maintaining health status in addition to cumulative food intake. The present study demonstrated that consumption of soluble dietary fiber inulin altered gut microbiota composition and induced a preference for a high-fat over a high-sugar diet. Existing literature has extensively discussed the benefits of dietary fiber on controlling food intake.

To date, this is the first study that examines the impact of dietary fiber inulin consumption on food preference between palatable high-fat and high-sugar diets. Dietary fiber inulin is known to control food intake and regulate one’s appetite, as commonly determined by total energy or cumulative food intake. Yet the effect of inulin on food preference has not been studied. Our results show that consuming dietary fiber inulin induces a preference for fat and suppresses the appetite for sugar in a mouse model. This observation suggests that previous dietary consumption substantially affects subsequent food choices. Nutrient-specific food selection operates on a positive feedback mechanism [29] in which pre-exposure to a certain macronutrient induces a preference for that macronutrient in mice. For example, the mice pre-exposed to fat self-selected fat, protein self-selected protein, and carbohydrates self-selected carbohydrates. A similar study demonstrated a contradictory result in which high pre-meal protein composition induced a low protein intake in subsequent meals with a reduction in total food intake [30]. Regardless, evidence demonstrates that previous meals substantially influence subsequent meals’ quantity and composition. Our results support the existing literature that the composition of the previous meal determines subsequent nutrient-specific food selection, even when the test diet compositions are not presented in the preceding meal. Preceding dietary composition’s effect on subsequent food choice was investigated more than three decades ago without consideration of gut microbiota as a component in the framework [30, 31]. Our study is one of the first that reinforces the effect of preceding dietary composition on subsequent food selection, considering gut microbiota as a factor.

Soluble dietary fiber consumption controls experimental animals’ energy intake and body weight [32, 33]; several studies have reported contradictory results. Some studies have reported that short-term inulin supplementation (12 days to 4 weeks) does not control body weight gain in experimental mice compared with controls [34–36]. In the present study, significant weight gain was detected in the inulin and control mice after the 2-week inulin intervention; no significant differences were detected in the post-dietary intervention body weights between the two groups. In particular, the mice in the present study were still undergoing their growth phase during the dietary intervention period (PD64-80), explaining the significant weight gain after dietary intervention. Several studies have reported that soluble dietary fiber increases metabolizable energy extraction from feed compared with insoluble dietary fiber, increasing body weight and leading to the development of the obese phenotype in mice [37, 38]. In the present study, we observed a tendency for higher weight gain in inulin mice than in the control. Determining the energy content of collected fecal pellets can be used to confirm this speculation in the future.

We compared nutrient-specific food selection in mice between two palatable test diets: a high-sugar diet (carbohydrate, 70.79%; fat, 11.82%; protein, 17.39%) and a high-fat diet (carbohydrate, 44.60%; fat, 40.38%; protein, 14.92%). Obesity results from a chronic energy imbalance between energy intake and expenditure, and food addiction exacerbates its development. The food addiction model is explained by neural activation in the reward circuitry in response to food cues and reduced activity in the inhibitory regions in response to food intake. Several reports have demonstrated that sugar is a food component that causes addictive behavior [39–43]. However, fat consumption does not elicit addictive behavior; overconsumption contributes to excessive weight gain from fat mass accumulation [39, 44, 45]. High-fat and high-sugar foods are ubiquitous in contemporary food environments and have differential downstream physiological effects when ingested. Food addiction is problematic because, unlike drugs and alcohol [4, 46], humans cannot eliminate food as it sustains survival. The present study assessed dietary fiber consumption’s effect on subsequent nutrient-specific food selection between palatable high-fat and high-sugar diets. Dietary fiber consumption regulates food and energy intake quantity and promotes a preference for fat over sugar, curbing food addiction development.

In the present study, three bacterial species positively correlated with fat preference: *Bacteroides acidifaciens, Bacteroides caecimuris*, and *Bacteroides xylanolyticus*, all belonging to the genus *Bacteroides*. These bacterial species were enriched in response to dietary fiber inulin consumption, consistent with several reports [47–49]. A recent study reported that the genus *Bacteroides*, particularly *Bacteroides uniformis CECT 7771*, is involved in reducing binge-eating behavior in rats, a process mediated by the serotonergic and dopaminergic pathways in the hypothalamus [50]. Another study revealed an inverse correlation between the relative abundance of *Bacteroides* and addiction-like eating behavior in obese women who underwent laparoscopic sleeve gastrectomy by reducing connectivity in the brain reward regions [51]. Combining the results from the present study illustrating that *Bacteroides* are positively correlated with a preference for a high-fat over a high-sugar diet, our findings imply *Bacteroides*’ protective role against addiction-like eating behavior in human and animal subjects. We aimed to explain the influence of the gut microbiota on food preference via the gut-brain axis. *Bacteroides spp*. were found to significantly induce short-chain fatty acids (SCFAs) production, particularly propionic acid, in the colon [48]. Sequentially, SCFAs induce serotonin in the gut [52], a neurotransmitter that suppresses carbohydrate intake [27, 31], providing a plausible explanation for the preference for a high-fat diet over a high-sugar diet observed in inulin mice in the present study. Moreover, the control mice in the present study exhibited a high abundance of *Bifidobacterium pseudolongum*, a bacterial species correlated with low serotonin concentration [53], which did not demonstrate a preference for either test diets. Therefore, we proposed a hypothesis: consumption of dietary fiber inulin induces the production of SCFAs and neurotransmitters that travel to the brain via the gut-brain axis to influence food preference (S7 Fig), in an attempt to demonstrate the underlying mechanism of food preference in inulin mice. Further studies involving fecal microbiota transplantation onto germ-free mice and measurement of candidate neurotransmitters are necessary to draw a causal relationship between gut microbiota composition and food preference.

## Conclusion

In summary, we demonstrated that dietary fiber consumption may induce a preference for a high-fat diet over a high-sugar diet, associated with the enrichment of gut bacteria *Bacteroides acidifaciens* (99.68%), *Bacteroides caecimuris* (99.37%), and *Bacteroides xylanolyticus* (92.28%). Our data on the impact of dietary fiber inulin on food preference may contribute to understanding behavioral consequences of dietary fiber ingestion on future food preferences through gut microbiota modulation.

## Acknowledgments

Yijia Liow designed and performed the experiments and wrote the original manuscript. Lena Takayasu financially supported and supervised the entire project and extensively revised the manuscript. Wataru Suda financially supported and provided important insights into the study design and revised the manuscript. Masahiro Umezaki provided insightful feedback to the manuscript. Itsuka Kamimura contributed to the design of the eating behavioral cage and the study design refinement. We thank Editage (www.editage.com) for English language editing.

## Supporting information

**S1 Fig. Customized eating behavioral testing cage**. The left- and right-chambers are the feeding chambers for mice to self-select between the test diets; the central area is the resting area for when the mice are not interacting with food. Marble balls are modifications made to the cage to prevent the mice from resting in the feeding chambers; stainless steel clips are used to secure the lid to prevent the mice from escape.

**S2 Fig. Individual and average fat preference score of individual mice across the 3-day eating behavioral test**. Inulin mice had a higher preference for the high-fat diet than did the control (P=0.0051).

**S3 Fig. Energy intake comparison between the inulin and control mice**. Fig 3a: No significant differences were detected in the energy intake of high-fat and high-sugar test diet energy intakes among the inulin and control mice (P=0.7 and 0.13, respectively). No significant differences were detected in the high-fat as well as high-sugar intake between the two groups (P=0.82 and 0.48, respectively). Fig 3b: Total energy intake combining high-fat and high-sugar test diets of the inulin and control mice; no significant differences were detected in the total energy intake between the two groups (P=0.66).

**S4 Fig. Mouse weight comparisons pre- and post-dietary intervention**. Mouse weight post-dietary intervention were higher in the inulin and control mice. Significant differences were detected in the mouse body weight gains before and after the dietary intervention in the inulin and control mice (P=0.0026 and 0.045, respectively). Mouse weight at baseline and post-intervention were not significantly different between the inulin and control mice (P=0.575 and 0.229, respectively).

**S5 Fig. Heatmap of the significantly different OTUs between the inulin and control mice post-dietary intervention (day 14) based on hierarchical clustering**. The colors on the heatmap reflect the OTU relative abundance z-score; red indicates OTUs high in relative abundance and blue indicates OTUs low in relative abundance.

**S6 Fig. Principal Coordinate Analysis (PCoA) plot based on weighted UniFrac distances for the gut microbiota composition in the inulin and control mice**. The legend indicates the chronological fecal sampling time points. The PCoA1 axis captures 28.96% of the total variation in the data and the PCoA2 axis captures 27.4% of the total variation in the data. PERMANOVA test reveals significant changes in the gut microbiota community structure in the inulin mice, starting from the third fecal sampling point (P=0.003).

**S7 Fig. Hypothesis underlying food preference induced by dietary fiber consumption**. Created with BioRender.com.

