## Supplementary figures and images for "Dietary fiber induces a fat preference associated with the gut microbiota"

### S1 Fig

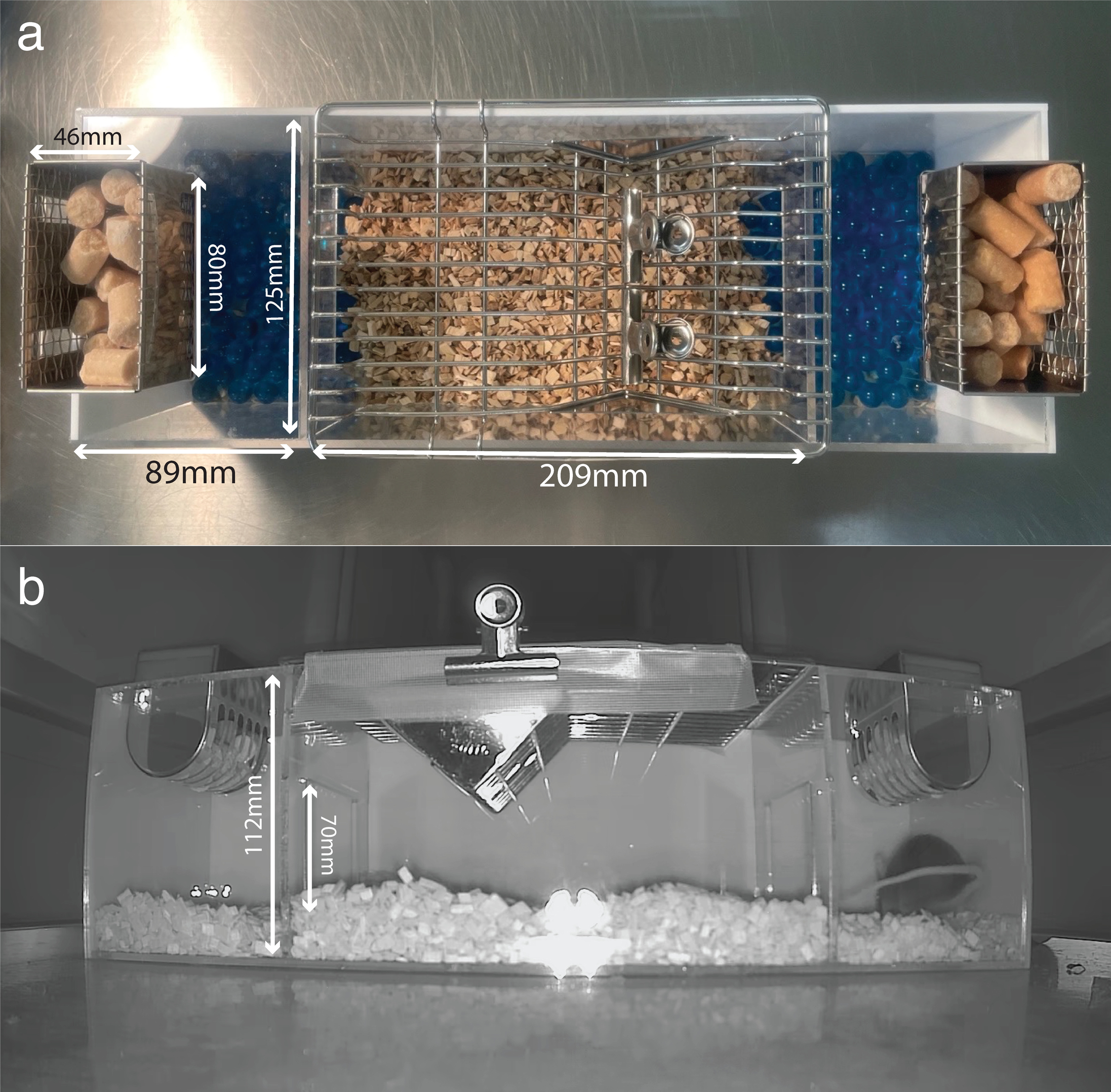

### S2 Fig

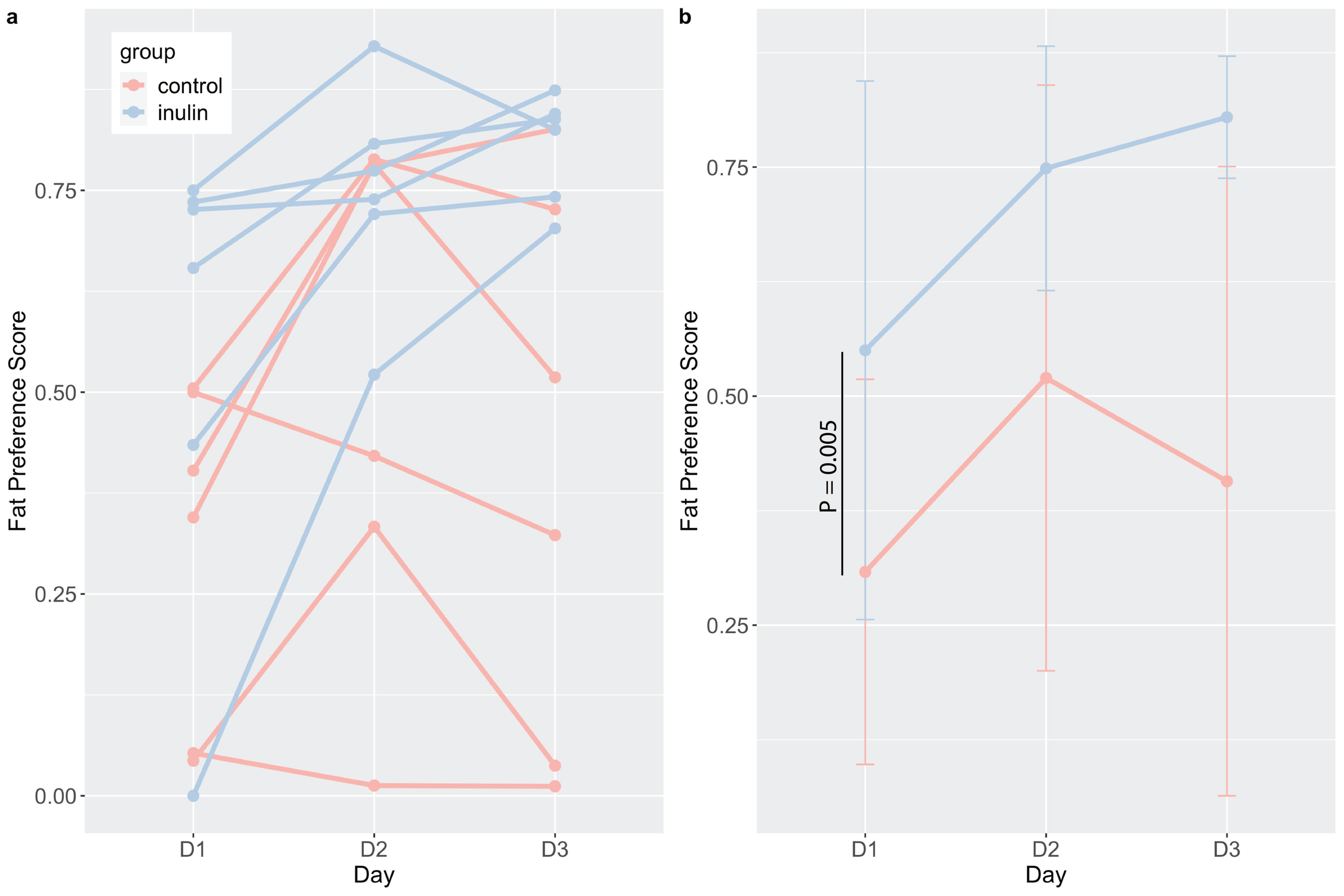

### S3 Fig

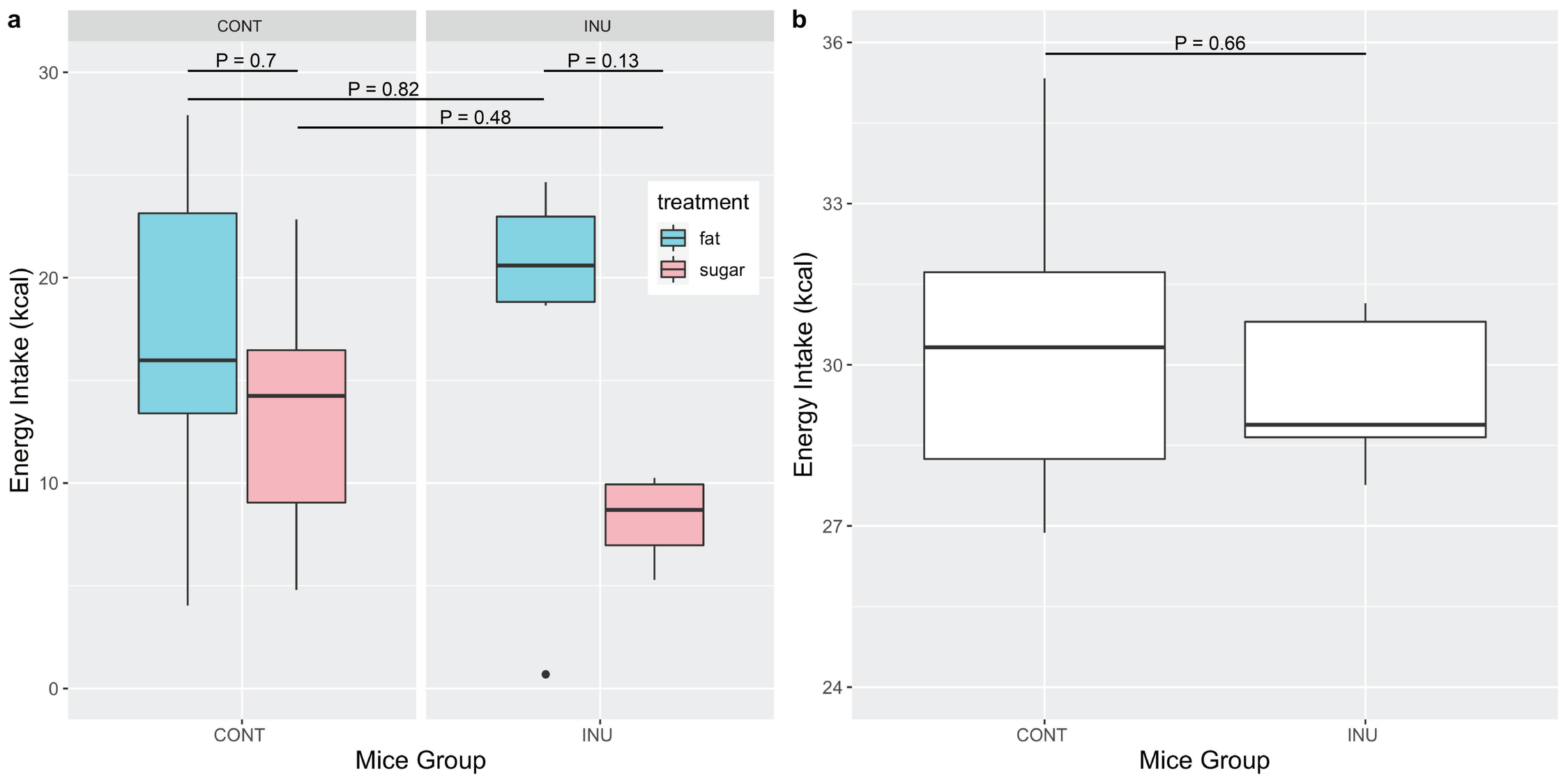

### S4 Fig

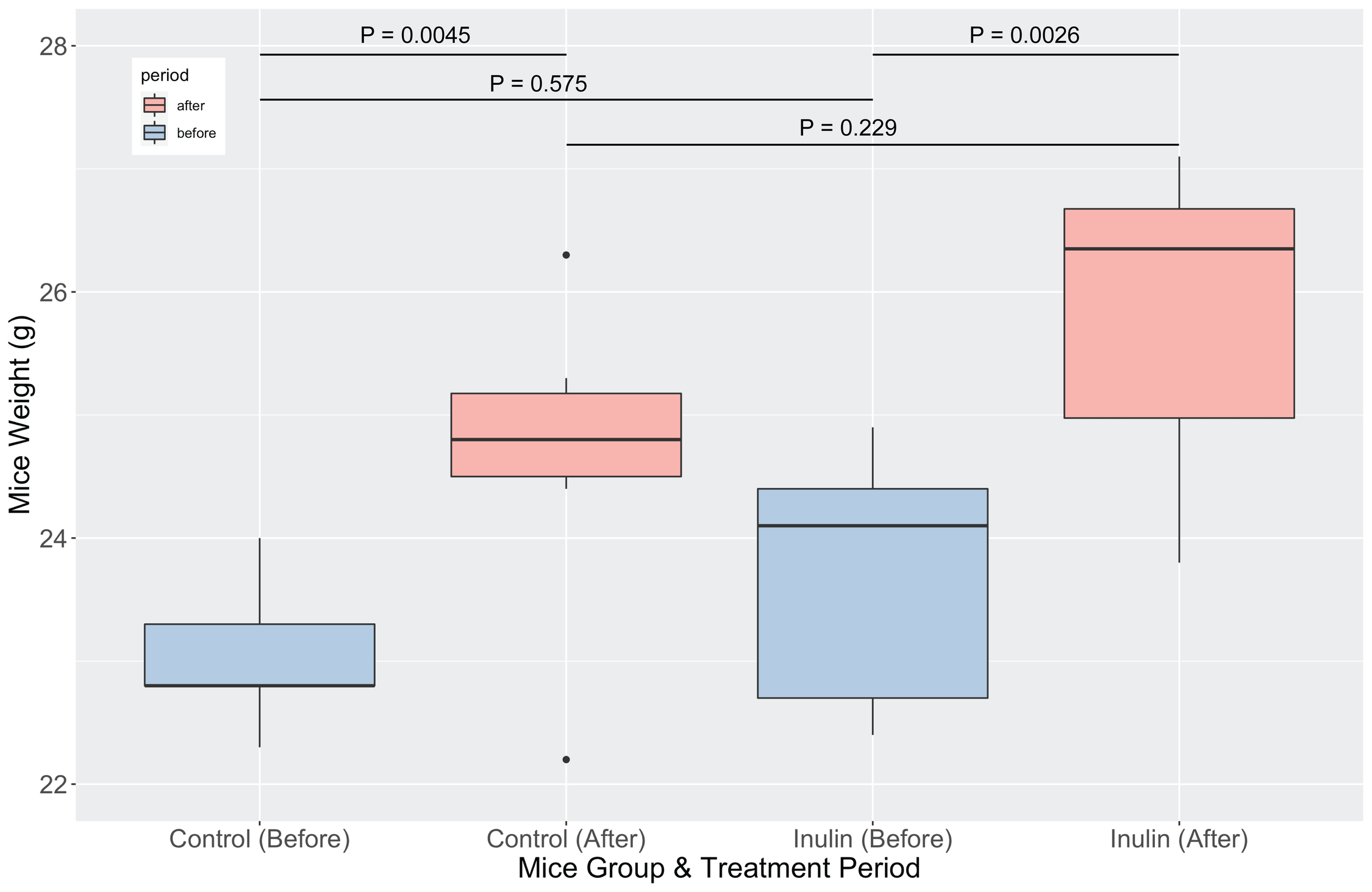

### S5 Fig

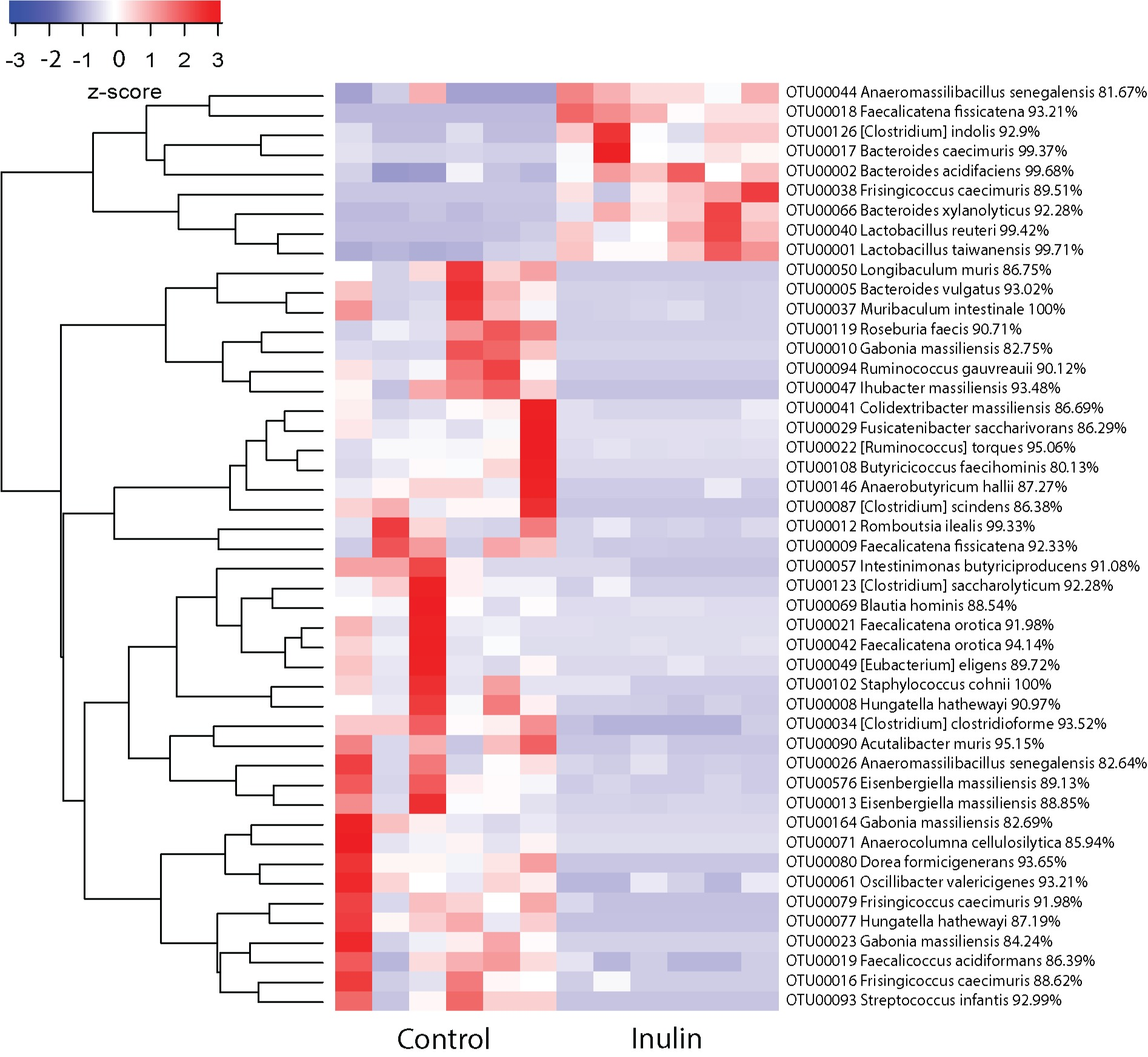

### S6 Fig

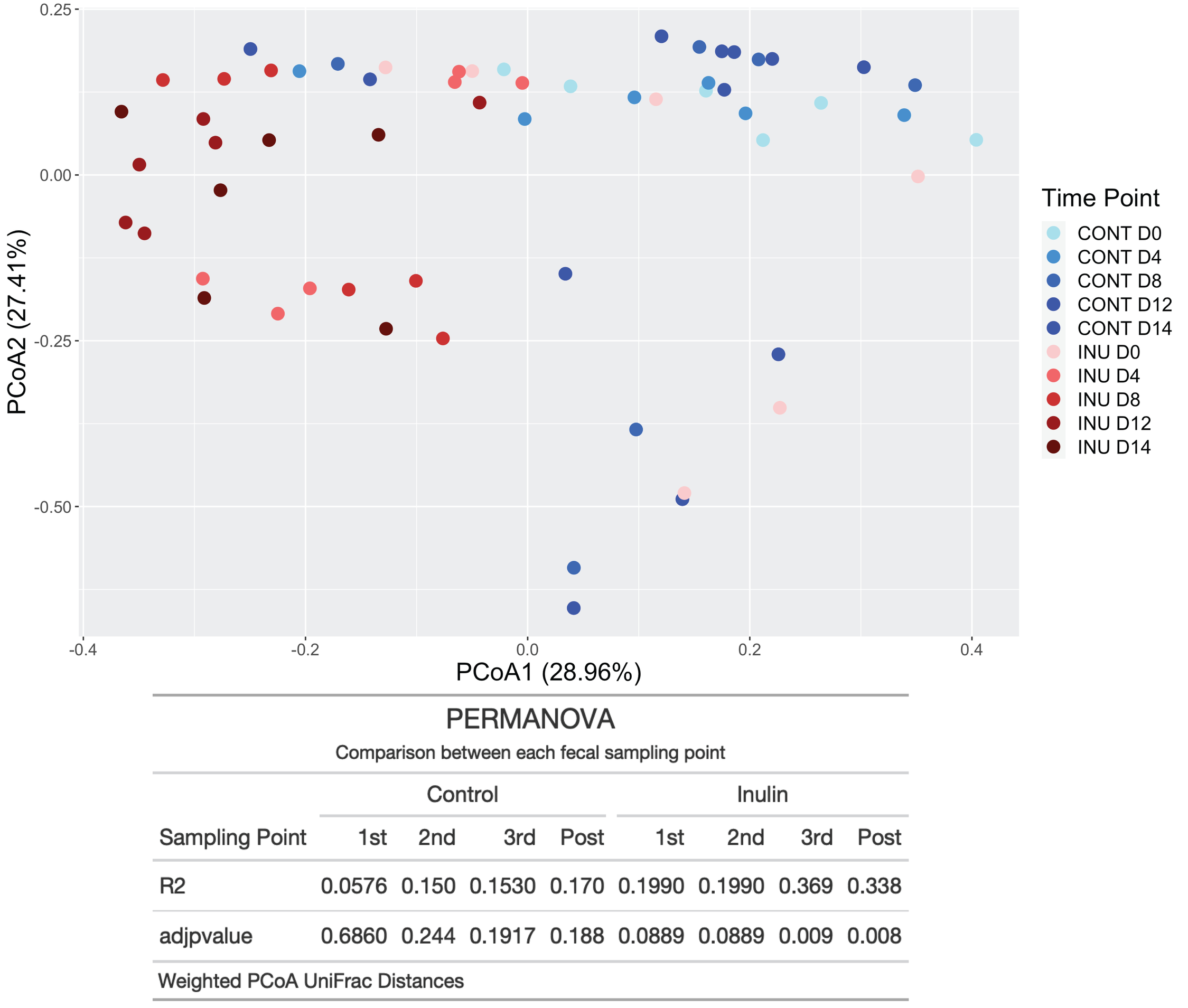

### S7 Fig

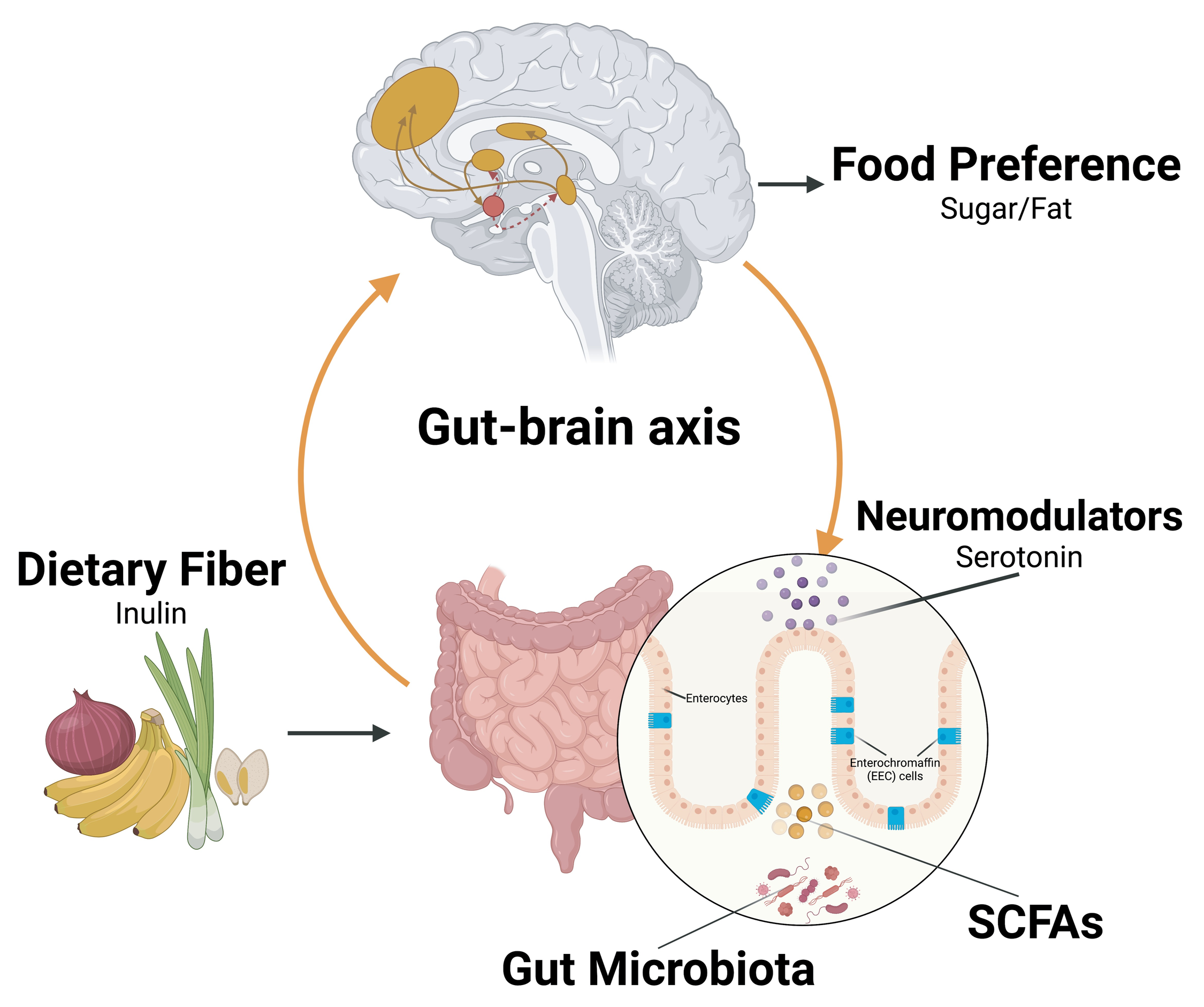
